# Nighttime-specific gene expression changes in suprachiasmatic nucleus and habenula are associated with resiliency to chronic social stress

**DOI:** 10.1101/2023.09.05.556324

**Authors:** Priyam Narain, Aleksa Petković, Marko Šušić, Salma Haniffa, Nizar Drou, Marc Arnoux, Mariam Anwar, Dipesh Chaudhury

## Abstract

The molecular mechanisms that link stress and circadian rhythms still remain unclear. The habenula (Hb) is a key brain region involved in regulating diverse types of emotion-related behaviours while the suprachiasmatic nucleus (SCN) is the body’s central clock. To investigate the effects of chronic social stress on transcription patterns, we performed gene expression analysis in the Hb and SCN of stress naive and stress exposed mice. Our analysis revealed a large number of differentially expressed genes and enrichment of synaptic and cell signalling pathways between resilient and stress-naïve mice at ZT16 in both the Hb and SCN. This transcriptomic signature was nighttime-specific and observed only in stress-resilient mice. In contrast, there were relatively few differences between the stress-susceptible and stress-naïve groups across timepoints. Our results reinforce the functional link between diurnal gene expression patterns and differential responses to stress, thereby highlighting the importance of temporal expression patterns in homeostatic stress responses.

## 1. Introduction

Mood disorders are complex heterogeneous disorders that affect cognition, emotional processing and basic physiological functioning (1). There is growing evidence of the close connection between circadian rhythms and mood regulation (2). For example, a majority of depressed patients show more severe symptoms in the morning than in the evening (3). Circadian rhythms are physiological, psychological and behavioural oscillations that follow an approximately 24-hour cycle. These internal rhythmic processes are maintained by multiple peripheral clocks throughout the body and are synchronised to a single central clock in the hypothalamus, the suprachiasmatic nucleus (SCN) in mammals (4,5). Circadian processes including diurnal rhythms in secretion of growth hormones, melatonin, daily core body temperature, and sleep/wake cycles are disrupted in some depressed patients (6–11). The SCN projects to several hypothalamic structures such as the paraventricular nucleus, subparaventricular zone, and dorsomedial hypothalamus that regulate these disrupted processes (6,12). The SCN also projects to regions that drive complex emotional behaviours such as the habenular complex (Hb), which is composed predominantly of lateral (LHb) and medial habenula (MHb) subregions (12,13). The direct and indirect projections of the SCN to mood-regulating areas of the brain are likely responsible for diurnal rhythms of pathological behaviours, such as anxious and depressive-like responses in mice (14,15).

The Hb relays information between the forebrain emotion-processing limbic system and the midbrain depression-associated aminergic centres (16,17). The Hb is heavily involved in mediating stress responses (18,19) and has also been associated with circadian modulation of depression (20). Pathophysiologically elevated firing in LHb neurons projecting to the serotonergic dorsal raphae nucleus (DRN) is associated with social avoidance in chronically stressed mice (21), while passive coping, a behaviour mirroring apathy and resignation in humans, has been associated with increased activity in numerous circuits including LHb-DRN projections (22–24), medial prefrontal cortex projections to LHb (25) and LHb projections to the ventral tegmental area (VTA) (26,27). The pharmacological inhibition of the LHb rescues depressive-like behaviour (23,28). In contrast, MHb lesion impairs voluntary exercise and induces despair-like and anhedonic behavioural phenotypes in mice, without affecting the circadian period and sleep architecture (29–31). Both LHb and MHb exhibit increased metabolism and structural changes in depressed patients (32,33). Despite its importance, the effects of stress on molecular pathways in the Hb remain underexplored.

The LHb exhibits an independent endogenous molecular clock that is synchronised by signals from the SCN (13,34–36). The SCN innervates the LHb directly (20,36,37) and indirectly through the dorsomedial and lateral hypothalamic areas (38–40). Moreover, the MHb sends direct projections into the LHb (41). Daily variation in the firing of both LHb and MHb neurons is driven by their functional molecular clock (20,35,42,43). Core clock genes are rhythmically expressed in the LHb in phase with SCN clock gene rhythms (13,44). Habenular clock robustly modulates complex behaviours, such as anxiety-like responses to stressors (45). Hence, depression-associated circadian disruption might be attributed to perturbations of circadian rhythms at the level of the SCN, the Hb or both.

Chronic social defeat stress (CSDS) is a robust and widely used chronic stress model that induces depression-like and PTSD-like symptoms in a subset of exposed rodents (46,47). Experimental mice exhibiting social avoidance are deemed “stress-susceptible”, while those that do not are “stress-resilient” (48). These two subpopulations show different behavioural, physiological and molecular profiles (21,46,49–51). This segregation allows investigations into underlying mechanisms that promote stress resilience (52,53). CSDS in rodents disrupts body temperature, heart rate, locomotor activity rhythms and sleep patterns where some processes are disrupted in all stress exposed mice and others only in the stress-susceptible group (51,54–56). We have shown that LHb cells projecting to the DRN of stress-susceptible mice exhibit blunted diurnal activity while stress-resilient and stress-naïve mice exhibit robust daily rhythm in activity (21). This observation suggests that abnormal diurnal rhythmic activity in LHb cells projecting to the DRN encodes susceptibility to stress. However, the temporal transcriptomic profiles of the Hb and SCN in the stress-resilient and stress-susceptible mice remains underexplored.

In this study, we investigated the effects of CSDS on the SCN and Hb transcriptome across the light/dark cycle. Mice exposed to CSDS were sampled at four timepoints: ZT4 (mid-inactive phase), ZT10 (transition to active phase), ZT16 (mid-active phase), and ZT22 (transition to inactive phase). We hypothesised that the greatest changes would be observed in the stress-susceptible mice in the Hb during the light phase. We expected relatively few differences between the resilient and stress-naïve mice, especially in the SCN. However, we found the largest transcriptomic changes at ZT16 in resilient relative to stress-naïve mice, in both SCN and Hb. Our observations also highlight the importance of measuring transcriptomic profiles across timepoints to determine the largest homeostatic change in response to stress.

## 2. Material and Methods

### 2.1. Animals and housing

Male C57BL/6J mice (8-10 weeks old; Jackson Laboratory) All mice were housed with *ad libitum* access to food and water at 20 ± 1 °C and 50 ± 10% humidity under a 12-hr light/dark cycle (7:00 am / ZT0: lights on; 7:00 pm / ZT12: lights off). All experimental procedures were conducted in accordance with the National Institute of Health Guide for Care and Use of Laboratory Animals and approved by the New York University Abu Dhabi Animal Care and Use Committee (IACUC Protocol: 19-0004).

### 2.2. Chronic social defeat stress

CSDS was performed as described previously (57). Experimental C57 mice were exposed to 10 minutes of physical stress followed by 24 hours of sensory stress in the cage of an unfamiliar CD1 aggressor mouse (Fig. S1A). CSDS was conducted daily over 15 days between ZT8 and ZT9. The control mice were housed in pairs in a separate room. All the mice were single-housed following the end of CSDS until sacrifice.

Social avoidance towards a novel non-aggressive CD1 mouse was measured in a two-stage social interaction (SI) test, as previously reported (57). The test was conducted between ZT7 and ZT10. TopScan video tracking system (CleverSys. Inc.) was used to automatically track mouse movement.

### 2.3. Tissue sampling and nuclei isolation

Mice were anaesthetized using isoflurane and sacrificed 48-72 hours after the SI test by cervical dislocation. Sampling was done at four timepoints (ZT4, ZT10, ZT16, ZT22) across the light-dark cycle (Fig.1A-B). Extractions were carried out on a cold sterile surface, and the extracted brains were flash-frozen by immersion into −60 °C isopentane (Sigma Aldrich M3263) for 20 seconds. The brains were kept on dry ice at −80°C until further processing. The flash-frozen brain samples were then sliced on a Leica CM1950 cryostat (250 µm thickness) (Fig. S1B). The nuclei were isolated with reference to the Mouse Brain Atlas (58). For both the SCN and Hb, bilateral punches from within each biological replicate were combined (Fig. S1B).

**Figure 1.**
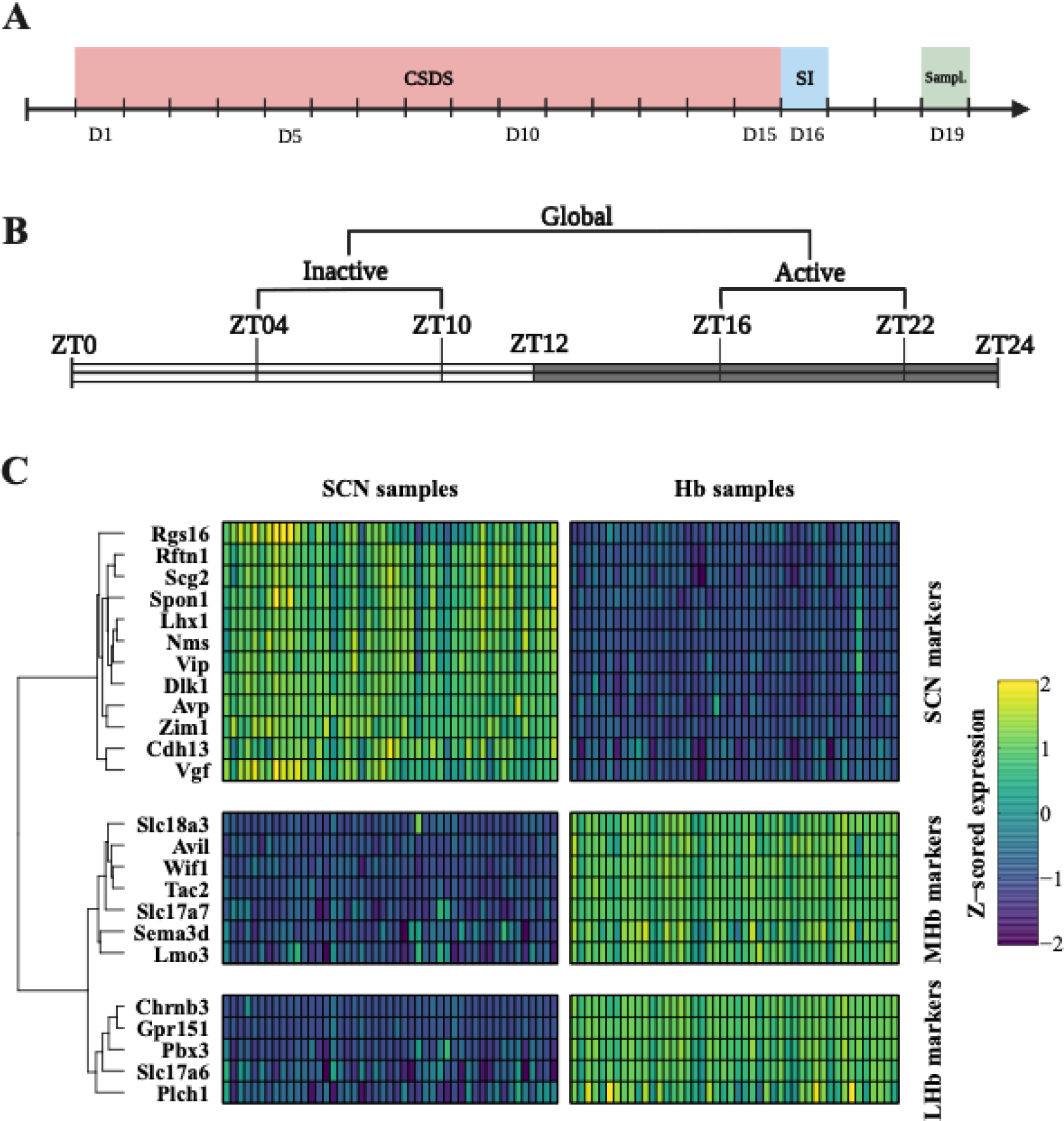
Experimental design and quality control. **(A)** Timeline for investigating the effect of stress on temporal changes in gene expression in stress-naïve and stress-exposed animals. **(B)** Sampling timepoints and emerging comparison levels from pooling the sampling timepoints. **(C)** Heatmap of canonical marker genes in the SCN and Hb. Normalized gene expression (variance stabilizing log2-transformed counts) across samples for reported marker genes specific to SCN, LHb and MHb (69,70).

### 2.4. RNA isolation, library preparation and RNA-sequencing

RNA was isolated from the punches using the Qiagen RNeasy Micro Kit, as per the manufacturer’s instructions. Isolated RNA was quantified and checked for quality on an Agilent Bioanalyzer instrument (Agilent Technologies) (Fig. S1C). Samples with RIN values above 7 were used for the library preparation. All the libraries were prepared together using the NEBNext® Ultra™ II RNA Library Prep Kit (New England BioLabs Inc.) as per the manufacturer’s protocol within one sitting. Sequencing was done on NovaSeq S1 flow cell (200 cycles) following Illumina’s recommendations.

### 2.5. Differential expression and pathway enrichment analysis

**Count preprocessing:** Raw FASTQ sequenced reads were first assessed for quality using FastQC (v0.11.5) (59). The reads were then passed through Trimmomatic (v0.36) for quality trimming (60) and processed with Fastp to remove poly-G tails ad Novaseq-specific artefacts (61). The reads were aligned to the mouse reference genome GRCm38.82 using HISAT2 with the default parameters (62). The resulting SAM alignments were then converted to BAM format and coordinate-sorted using SAMtools (v1.3.1) (63). Raw counts were generated by passing the sorted alignment files through HTSeq-count (v0.6.1p1) (64). Concurrently, the sorted alignments were processed through Stringtie (v1.3.0) for transcriptome quantification (65). Finally, Qualimap (v2.2.2) was used to generate RNAseq-specific QC metrics per sample (66). All of the methods were executed using predefined YAML workflows using the BioSAILs workflow management system (67).

**Differential expression**: Differential expression testing was performed in R package DESeq2 (v1.40.2) using multiple Wald tests (68). To control for the false discovery rate, *p* values were adjusted using the Benjamini–Hochberg procedure. To obtain a better estimate of expression changes in low count genes, we performed log2-fold change (LFC) shrinkage using the *apeglm* method. We used an adjusted *p* value of 0.05 (5% FDR) and a log2-fold change (LFC) of 0.5 to identify differentially expressed genes (DEGs). Additional quality control was conducted to account for technical errors during sequencing. Samples within the same behavioural phenotype from ZT4 and ZT10 timepoints were computationally pooled together to provide an estimate of the ‘daytime’ expression, while ZT16 and ZT22 timepoint samples were pooled to provide an estimate of the ‘nighttime’ expression for each phenotype (Fig.1B). Pooling all the samples within the same behavioural phenotype provided an estimate of time-invariant ‘global’ expression corresponding to average expression throughout the day. Differential expression between the behavioural phenotypes was carried out at each of these levels. All the sample identities were assessed using previously published regional markers (Fig.1C; Fig. S2) (69,70).

**Figure 2.**
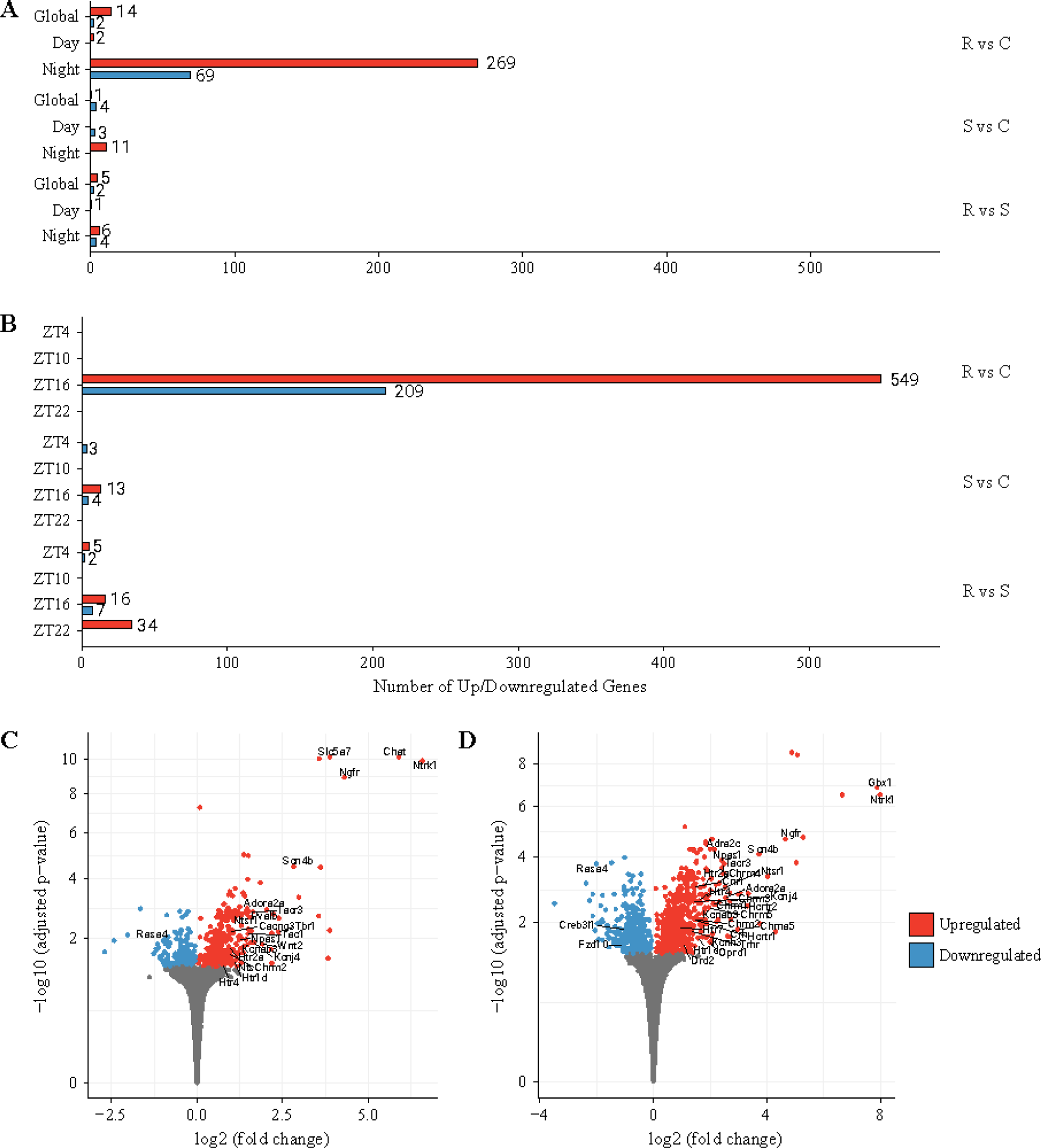
Differentially expressed genes (DEGs) at different time point levels across phenotypes in the SCN. **(A)** Number of DEGs at pooled timepoints. At the global comparison level, there were 16, 5 and 7 DEGs (FDR < 0.05 and |log2FC| > 0.5) in resilient-to-control, susceptible-to-control and resilient-to-susceptible comparisons, respectively. Daytime comparison revealed 2, 3 and 1 DEGs for resilient-to-control, susceptible-to-control and susceptible-to-resilient comparisons, respectively. Night time comparison revealed 338, 11 and 10 DEGs for resilient-to-control, susceptible-to-control and susceptible-to-resilient comparisons, respectively. **(B)** Number of DEGs at specific timepoints. At ZT4, there were 0, 3 and 7 DEGs for resilient-to-control, susceptible-to-control and resilient-to-susceptible comparisons. At ZT10, there were no DEGs for any phenotypic comparison. At ZT16, there were 758, 17 and 23 DEGs for resilient-to-control, susceptible-to-control and resilient-to-susceptible comparisons, respectively. At ZT22, there were 0, 0 and 34 DEGs for resilient-to-control, susceptible-to-control and resilient-to-susceptible, respectively. **(C)** Nighttime comparison. Volcano plot showing the DEGs in resilient mice relative to controls at nighttime comparison level in SCN. **(D)** ZT16 comparison. Volcano plot showing the significant differentially expressed genes in resilient mice relative to controls at ZT16 in SCN. Red and blue points represent significantly up-regulated and down-regulated genes, respectively.

**Pathway analysis**: Enrichment analysis was conducted in the Ingenuity Pathway Analysis (IPA, Qiagen) on all genes that had passed the 5% FDR significance criterion and was conducted only in those comparisons where the number of differentially expressed genes (DEGs) was higher than 20. Pathways were considered significantly enriched if –log(B-H *p-*value) was greater than 1.3. Based on the known roles of different genes and their relationship in the functionally-predefined pathways, IPA calculates the *z* score which represents the predicted direction of change of the pathway (activation: *z* > 0; inhibition: *z* < 0). The analysis included only the pathways associated with the mouse nervous system, neurons, astrocytes, other tissues and unspecified cell types.

**Gene ontology:** We visualised the associated DEGs and the top 5 IPA pathways from the resilient-to-control comparisons at ZT16 for both nuclei using gene-concept networks. Gene-concept networks represent the relative coordinates of the enriched pathways based on the number and differential expression of the associated DEGs. Therefore, it is possible to identify overlapping DEGs that have the largest contribution to the enrichment of the top 5 activated pathways.

**Detection of rhythmic genes:** Log-normalised count values were used to determine the rhythmic expression profile of core clock genes across all three behavioural phenotypes using the non-parametric JTK method for identification of oscillations in genome-scale datasets (71). JTK method was run in the R-implemented version of the ECHO (v.4.0) tool (72).

## 3. Results

### 3.1. CSDS induces social avoidance in a subset of stress-exposed mice

To investigate the effects of chronic stress on SCN and Hb transcriptome, we exposed mice to 15 days of CSDS. Following CSDS, mice were classified as stress-resilient or stress-susceptible (Fig. S3A-C) (Golden et al., 2011). Stress-naïve and stress-resilient mice exhibited significantly larger social interaction values than the stress-susceptible mice (Fig. S3A-C). Resilient and susceptible mice exhibited decreased movement and velocity relative to the stress-naïve mice (Fig.S3D-E).

### 3.2. Stress-resiliency is associated with large gene upregulation in SCN and Hb at ZT16

In order to investigate differential expression between the behavioural phenotypes at different timepoints, we sampled SCN and Hb at ZT4, ZT10, ZT16 and ZT22. We observed the highest number of DEGs between resilient and stress-naïve mice at all levels of comparison in the SCN and Hb. The majority of these DEGs were observed in the resilient mice relative to the stress-naïve mice.

In the SCN (Fig. 2A-B), at global (average gene expression throughout the day) and daytime (average daytime gene expression) comparisons only a few genes were differentially expressed between any of the groups. Comparison of gene expression patterns at ZT4 and ZT10 timepoints did not reveal substantial differences between the phenotypes. However, at nighttime, we found 338 DEGs in resilient relative to stress-naïve mice, out of which 269 were upregulated (Fig.2A,C). The nighttime differences in the SCN is predominantly from the differential expression at ZT16 where we found 1058 DEGs in resilient relative to stress-naïve mice, out of which 549 were upregulated (Fig.2B,E). At ZT22, we observed 34 DEGs in resilient relative to susceptible mice. However, other phenotypic comparisons at ZT22 did not reveal any major differential expression patterns. These results suggest that the largest changes in stress-induced gene expression observed in the SCN occur in resilient mice during the nighttime at ZT16.

In the Hb (Fig.3A-B) differential expression patterns between the phenotypes in terms of number and direction of DEGs were similar to those observed in the SCN. Global comparisons in the Hb revealed 68 upregulated DEGs in the resilient relative to stress-naïve mice (Fig.3A,C). Gene expression patterns did not robustly differ between any of the three behavioural phenotypes during the daytime. Specifically, ZT4 and ZT10 timepoint comparisons did not yield any discernible differences in gene expression patterns (Fig. 3B). However, nighttime comparison between the behavioural phenotypes revealed 55 upregulated DEGs in resilient relative to stress-naïve mice (Fig.3A,D). We found that the large differences in global and nighttime gene expression profiles are mainly due to the large numbers of upregulated genes at ZT16 in stress-resilient relative to control and susceptible mice (Fig.3A-B). Specifically, 143 DEGs were identified in the resilient relative to stress-naïve mice at ZT16, out of which 129 genes were upregulated (Fig. 3B,E). We observed 78 upregulated DEGs in resilient mice compared to stress-susceptible mice, but there were no differences between the susceptible and control mice (Fig. 3B). No robust differences were observed at ZT22 between any of the behavioural phenotypes (Fig.3B). Hence, stress-induced differences in Hb gene expression patterns in the resilient mice, both at global and nighttime analysis, are driven by changes at the ZT16 timepoint.

**Figure 3.**
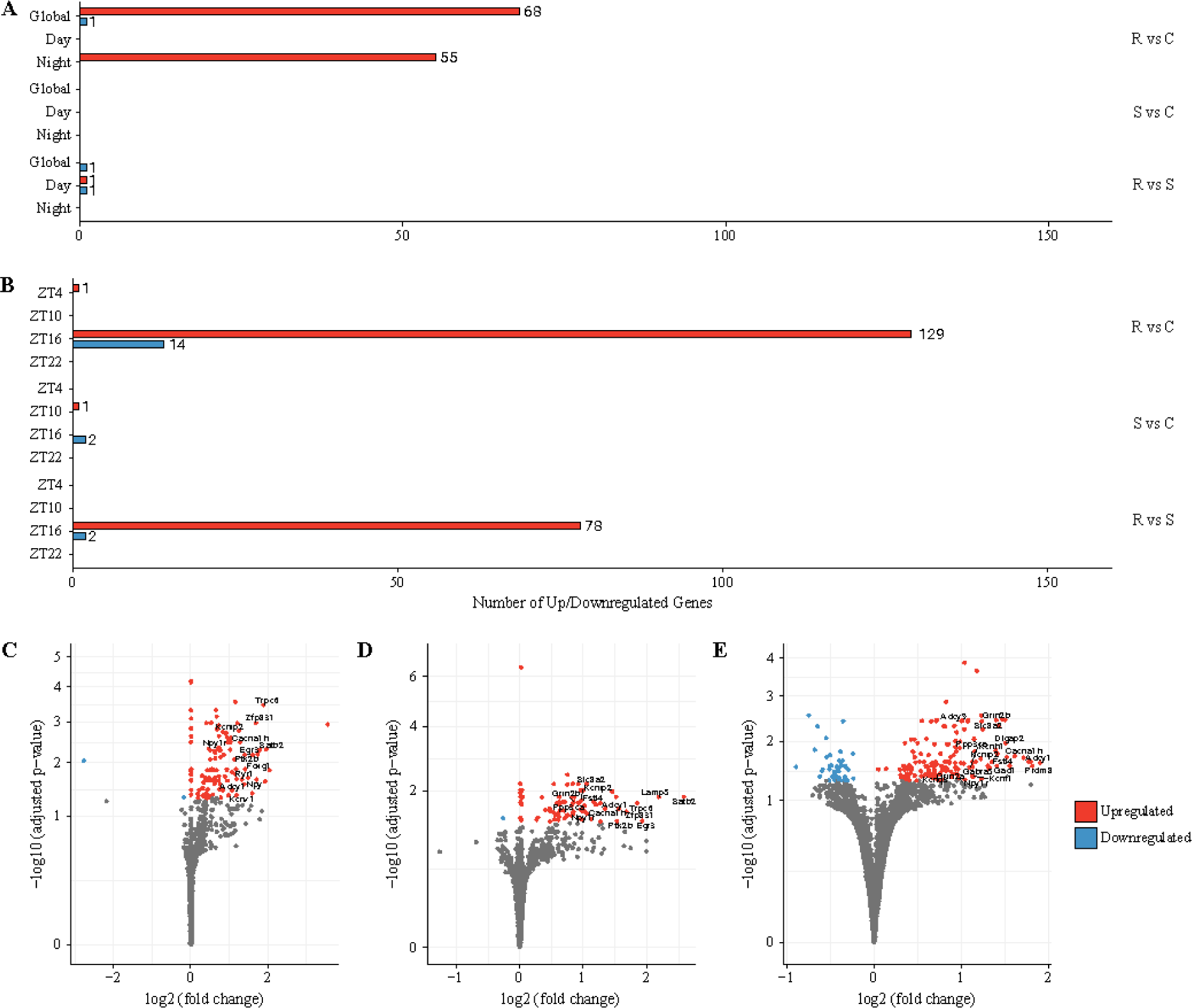
Differentially expressed genes (DEGs) at different time point levels across phenotypes in the Hb. **(A)** Number of DEGs at pooled timepoints. At the global comparison level, there were 69, 0 and 1 DEGs for resilient-to-control, susceptible-to-control and resilient-to-susceptible comparisons, respectively. At the daytime comparison level, there were 0, 0 and 2 DEGs for resilient-to-control, susceptible-to-control and susceptible-to-resilient comparisons, respectively. At the nighttime comparison level, there were 55, 0 and 0 DEGs for resilient-to-control, susceptible-to-control and susceptible-to-resilient comparisons, respectively. **(B)** Number of DEGs at specific sampled timepoints. At ZT4, there were 1, 0 and 0 DEGs for resilient-to-control, susceptible-to-control and resilient-to-susceptible comparisons. At ZT10, there were no DEGs for resilient-to-control and resilient-to-susceptible comparisons, and only 1 DEG in susceptible-to-control comparison. At ZT16, there were 143, 2 and 80 DEGs for resilient-to-control, susceptible-to-control and resilient-to-susceptible comparisons, respectively. At ZT22, there were no DEGs in any of the phenotypic comparisons. **(C)** Global comparison. Volcano plot showing the DEGs in resilient mice relative to controls at global comparison level in Hb. DEGs were labeled based on |log2FC| > 1.0 and biological relevance. **(D)** Nighttime comparison. Volcano plot showing the DEGs in resilient mice relative to controls at nighttime comparison level in Hb. **(E)** ZT16 comparison. Volcano plot showing the DEGs in resilient mice relative to controls at ZT16 timepoint in Hb. Red and blue points represent significantly upregulated and downregulated genes, respectively.

Overall, these results show robust transcriptomic differences in resilient relative to stress-naïve mice in both the Hb and SCN. The largest observed differences occur at ZT16 in both nuclei, with SCN showing larger transcriptional response than the Hb in terms of number of DEGs and their LFC. Susceptible mice did not show significant differential expression patterns relative to either the resilient or control mice. Taken together adaptive homeostatic responses in the resilient mice at ZT16 leads to greater DEG in resilient mice.

### 3.3. Stress resilience is associated with enrichment of signalling and plasticity pathways in SCN at ZT16

To better understand the molecular pathways and functions of DEGs between the resilient and stress-naïve mice, we performed pathway enrichment analysis in the SCN at nighttime and at ZT16 and found a number of enriched pathways (Table 1, Fig.4). At ZT16, the enriched pathways included an expanded list of those observed from the nighttime comparison (Table 1, Fig.4A-B). All the enriched in the resilient relative to stress-naïve mice are divided into two broad functional groups: signalling and plasticity-related pathways (Fig.4A). Furthermore, at ZT16, circadian rhythm signalling pathway was significantly enriched in the resilient mice. Both signalling and plasticity-related pathways in the resilient mice were enriched by common DEGs, a subset of which is shown in Table 2. Many upregulated DEGs were associated with relatively few pathways (Fig.4C). These results suggest that stress resilience is strongly associated with upregulation of relatively few genes in SCN at ZT16 which have broad impact on signalling and plasticity regulation.

**Figure 4.**
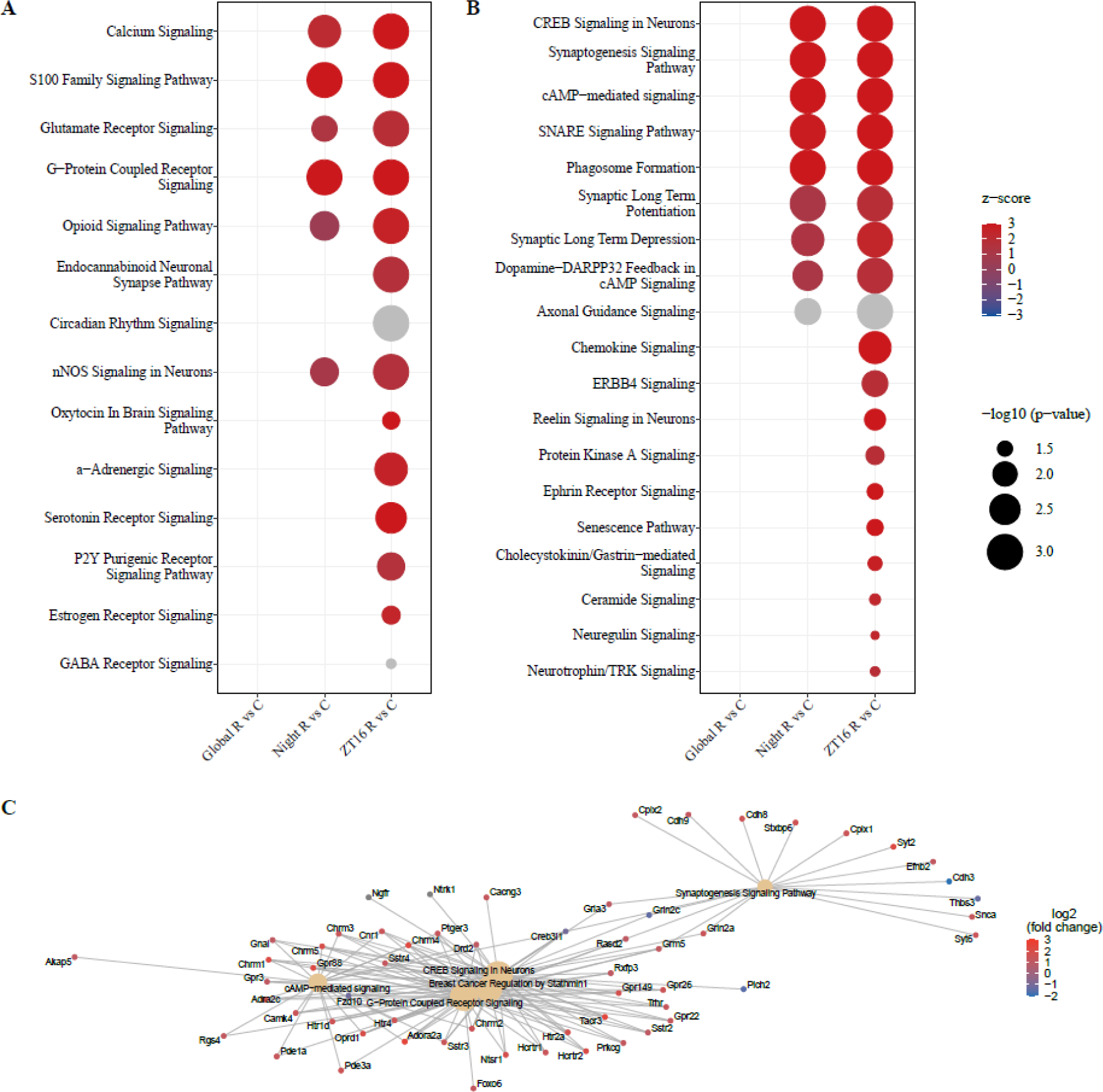
Enriched pathways in the SCN of resilient mice relative to controls. The activation pattern of enriched biological pathways in the SCN. All lists are sorted by adjusted p-value. Red denotes activation (*z* > 0), blue denotes inhibition (*z* < 0), white denotes no net effect (*z* = 0), and grey denotes unpredicted activity. **(A)** Enriched signaling pathways. **(B)** Enriched plasticity-associated pathways. **(C)** Gene-concept networks of the 5 most enriched IPA pathways at ZT16 between resilient and control phenotypes in the SCN.

**Table 1.** IPA-predicted *z* score of enriched pathways between the stress-resilient and stress-naive mice in the SCN. Only relevant pathways with |*z|* > 2.0 in any of the comparisons are shown.

| Enriched Pathway | Nighttime<br>R vs C | ZT16<br>R vs C |
| --- | --- | --- |
| CREB signaling in neurons | 5 | 6.3 |
| Phagosome formation | 4.7 | 6 |
| S100 family signaling | 4.6 | 5.2 |
| G-protein coupled receptor signaling | 3.9 | 5.2 |
| Synaptogenesis signaling | 3.8 | 5.1 |
| Serotonin receptor signaling |  | 3.9 |
| cAMP-mediated signaling | 3.5 | 3.8 |
| Neurovascular coupling signaling | 3.0 | 3.8 |
| SNARE signaling | 2.8 | 3.5 |
| Chemokine signaling |  | 3.5 |
| Reelin signaling in neurons |  | 3.4 |
| Oxytocin in brain signaling |  | 3.3 |
| Calcium signaling | 2.1 | 3.1 |
| Ephrin receptor signaling |  | 3.1 |
| Cholecystokinin/gastrin-mediated signaling |  | 2.7 |
| ERBB signaling |  | 2.7 |
| Opioid signaling |  | 2.6 |
| α-adrenergic signaling | | 2.5 |
| Synaptic long-term depression |  | 2.4 |
| Estrogen receptor signaling |  | 2.4 |
| Neuregulin signaling |  | 2.3 |

**Table 2.** DEGs associated with 4 or more enriched IPA pathways in resilient relative to stress-naive mice at ZT16. All presented genes have |LFC| > 1.0. Genes colored red are upregulated, while genes colored blue are downregulated.

| Gene Functional Group | SCN | Hb |
| --- | --- | --- |
| Ion channels | *Cacng3, Gabra1, Grin2a, Grin2c, Kcnj4* | *Cacna1h, Grin2a* |
| Receptors | *Adora2a, Adra2c, Chrm1-5, Cnr1, Drd2, Fzd10, Gnal, Grm5, Hctr1-2, Hrh2, Htr1d, Htr2a, Htr4, Ngfr, Ntsr1, Oprd1, Ptger3, Rxfp3, Sstr 2-4, Tacr3, Trhr* |  |
| Enzymes | *Camk4, Ngef, Nos1, Prkcg* | *Camk1d, Gad1, Ppp3ca* |
| Other | *Creb3l1, Rasd2, Rgs2, Rgs7* | *Adcy1, Adcy9* |

### 3.4. Stress resilience is associated with enrichment of synaptogenesis and calcium signalling pathways in Hb at ZT16

In the Hb, we explored pathway enrichment at global, nighttime and ZT16 comparisons between resilient and stress-naïve mice (Fig.5). Fewer pathways were as highly enriched and activated in the Hb of the resilient mice at ZT16 compared to the SCN (Table 3, Fig.5A-B). The most enriched pathways in this comparison are associated with mechanisms of synaptic plasticity (Table 3, Fig. 5B). Similarly to the SCN, circadian rhythm signalling pathway was enriched in the resilient mice at global, nighttime and ZT16 comparisons. Relatively few DEGs were associated with a high number of enriched pathways in the Hb of resilient mice (Table 2). Furthermore, the most enriched pathways were associated with only handful of genes (Fig. 5C). These results suggest that stress resilience is associated with upregulation of plasticity-associated genes in Hb at ZT16. Although both the SCN and Hb exhibited robust changes specifically at ZT16, SCN shows larger and functionally more diverse transcriptomic response to stress than the Hb.

**Figure 5.**
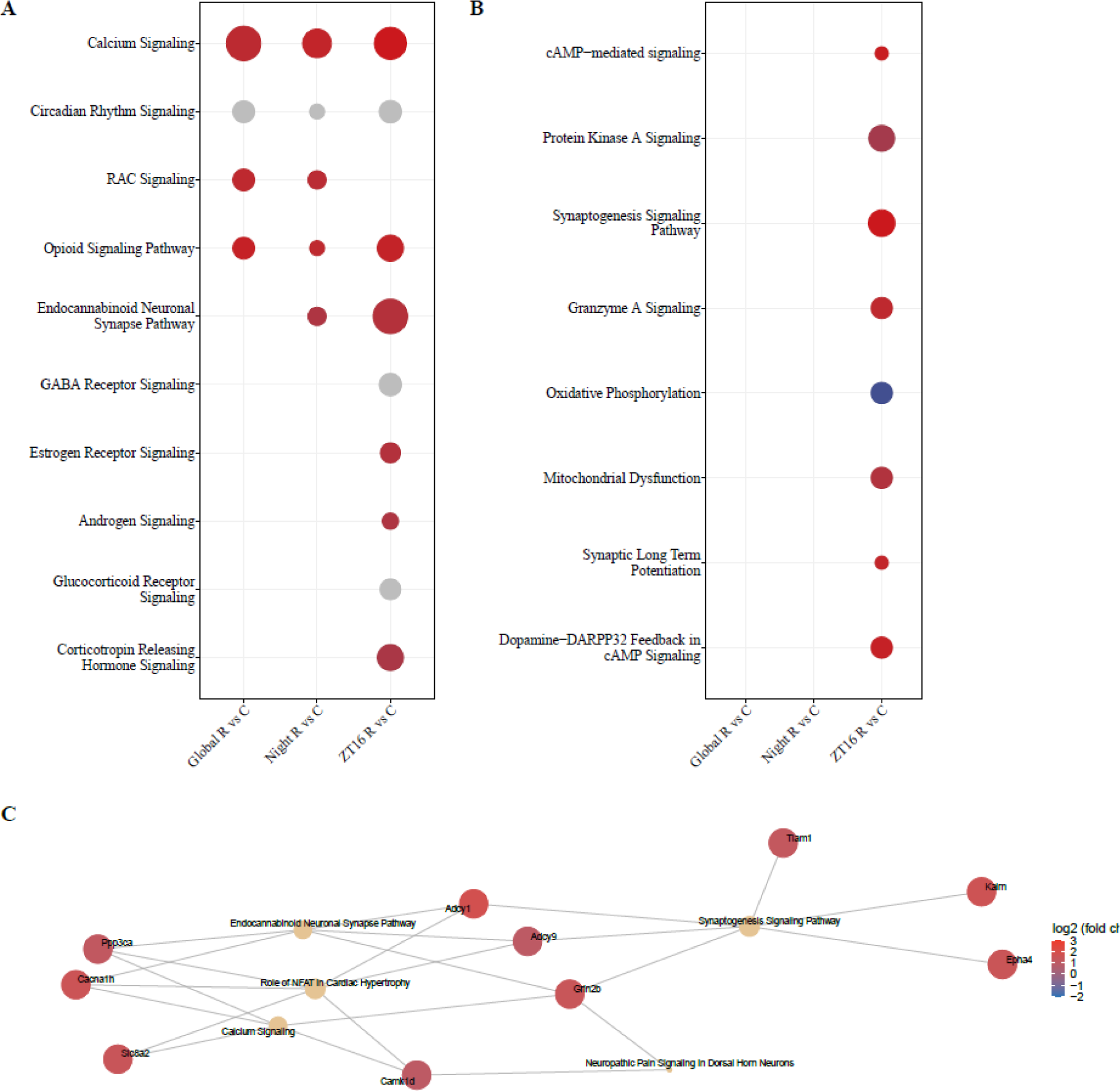
Enriched pathways in the Hb of resilient mice relative to controls. The activation pattern of enriched biological pathways in the Hb. All lists are sorted by adjusted p-value. Red denotes activation (*z* > 0), blue denotes inhibition (*z* < 0), white denotes no net effect (*z* = 0), and grey denotes unpredicted activity. **(A)** Enriched signaling pathways. **(B)** Enriched plasticity-associated pathways. **(C)** Gene-concept networks of 5 most enriched IPA pathways at ZT16 between resilient and control phenotypes in the Hb.

**Table 3.**
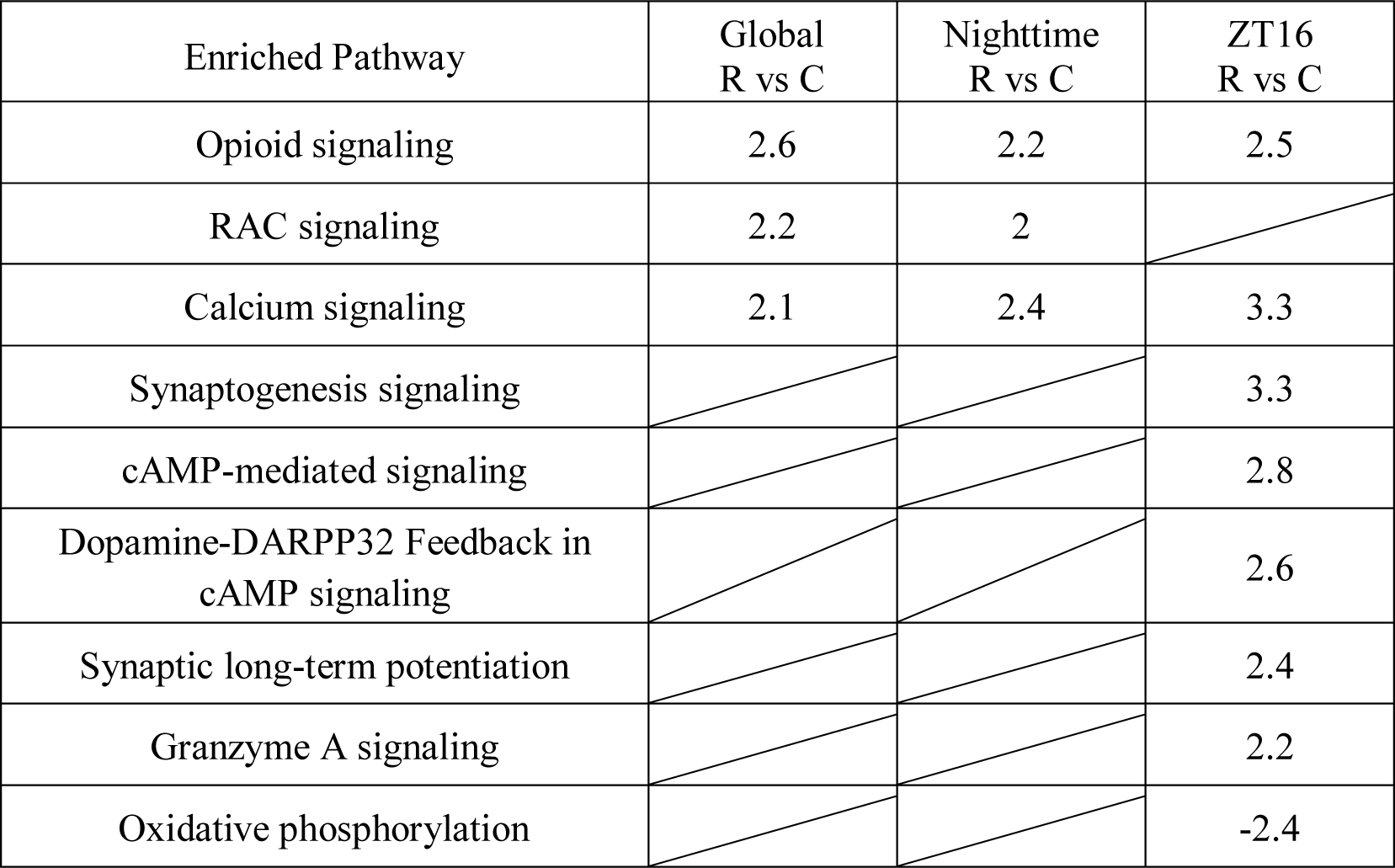
IPA-predicted *z* score of enriched pathways between the stress-resilient and stress-naive mice in the Hb. Only relevant pathways with |*z|* > 2.0 in any of the comparisons are shown.

| Enriched Pathway | Global<br>R vs C | Nighttime<br>R vs C | ZT16<br>R vs C |
| --- | --- | --- | --- |
| Opioid signaling | 2.6 | 2.2 | 2.5 |
| RAC signaling | 2.2 | 2 |  |
| Calcium signaling | 2.1 | 2.4 | 3.3 |
| Synaptogenesis signaling |  |  | 3.3 |
| cAMP-mediated signaling |  |  | 2.8 |
| Dopamine-DARPP32 Feedback in<br>cAMP signaling |  |  | 2.6 |
| Synaptic long-term potentiation |  |  | 2.4 |
| Granzyme A signaling |  |  | 2.2 |
| Oxidative phosphorylation |  |  | -2.4 |

### 3.5. Stress induces arrhythmicity of *Per1* and *Artnl* in SCN of stress exposed mice

Given the enrichment of circadian rhythmicity signalling pathway, we explored if chronic stress affected the rhythmicity of core clock genes in the SCN and Hb. To this end, we performed JTK analysis and found as reported previously rhythmic expression of the core clock genes (*Per1*, *Per2*, *Per3*, *Cry1*, and *Arntl*) in the SCN (p_adj_ <0.05) of stress-naïve mice (Fig.6A-B). Resilient and susceptible mice exhibited similar rhythmic expression profiles of the core clock genes in the SCN as stress-naïve mice, with the exception of *Arntl* and *Per1*. *Arntl* was arrhythmic in both stress-resilient and stress-susceptible mice, while *Per1* was arrhythmic in the SCN of stress-susceptible mice. In the Hb the expression of core clock genes were rhythmic in all three behavioural phenotypes, with the exception of *Per1, Per3,* and *Cry1* which were arrhythmic in the Hb of stress-naïve mice (Fig.6A-B).

**Figure 6.**
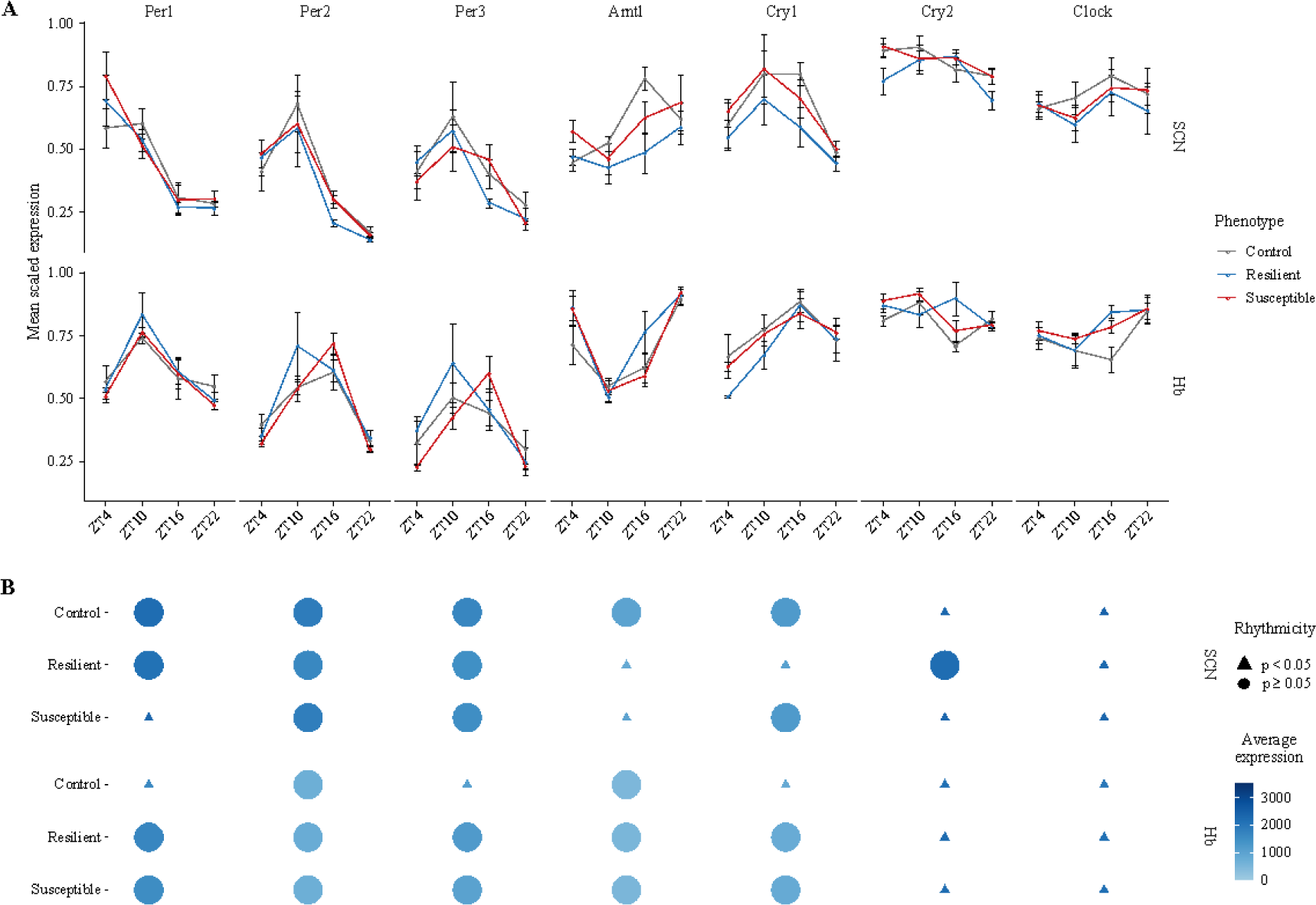
Clock gene rhythmicity. **(A)** Temporal expression of core clock genes in the SCN and Hb of control, resilient and susceptible mice. All the clock genes show previously reported temporal expression in the SCN of control mice. **(B)** Dotplot showing rhythmic expression of clock genes in SCN and Hb. **SCN**: *Per1* and *Cry1* are arhythmic in susceptible and resilient mice respectively. *Arntl* is arrhythmic in both resilient and susceptible mice. **Hb**: *Per1*, *Per3* and *Cry1* are arrhythmic in the control mice, while being rhythmic in both the resilient and susceptible mice.

## 4. Discussion

### Stress resilience and homeostatic response

The close relationship between circadian rhythms and mood disorders warranted a systemic investigation of transcriptomic changes in the principal clock (SCN) and brain regions that modulate mood. We focused on the Hb because it is a semi-autonomous oscillator with strong association with stress and mood disorders (20,45,73). We found that chronic stress elicited a robust change in the gene expression in the SCN and to a lesser degree in the Hb (Fig.2 & 3). This transcriptomic signature was specific to resilient mice and was strongest at ZT16. In contrast, comparing susceptible and stress-naïve mice did not reveal strong differential expression patterns. Large differential upregulation in resilient mice was associated with enrichment and predicted activation of signalling and plasticity-related pathways in both nuclei (Fig.4 & 5). Resilience is an active homeostatic process and involves transcriptional changes in a diffuse network of interconnected brain regions (74,75). The current understanding is that chronic stress imposes an adaptive challenge that requires behavioural flexibility and physiological homeostatic buffering (76). The inability to elicit such a response leads to failure in stress-coping and strongly contributes to associated stress-induced pathologies (77,78). Resilient mice do not show behavioural deficits likely because of large transcriptomic changes such as ones observed in our study. Early studies found that resilient mice exhibited large and unique upregulation of neurotrophic factors and stress-responsive genes in the prefrontal cortex (46). Others found strong gene upregulation in resilient mice relative to control and susceptible mice in midbrain regions such as the nucleus accumbens and VTA (79–82). Our findings that large transcriptomic changes occur in resilient mice at specific night timepoint (ZT16) add a significant temporal dimension to our understanding of molecular mechanisms that drive stress-resilience. Our data, along with previously published physiological and circuit studies, suggest that temporal patterns of regional activity may play a crucial role in driving a particular behavioural phenotype. For example, stress-susceptible mice exhibit elevated firing in the LHb during the day relative to stress-resilient and stress-naïve mice, suggesting a diurnal disruption at the level of LHb (21). This increased burst firing in the LHb is in part driven by signals from other stress-response regions (21,83). However, the functional consequences of large transcriptomic changes in resilient mice are likely responsible for differences in circuit-specific neural activity between resilient and susceptible mice. For example, VTA dopaminergic (DA) cells in susceptible, but not resilient, mice exhibit elevated firing (46,84,85). The pathophysiologically elevated firing is counteracted in resilient mice by compensatory mechanisms that are absent in susceptible mice (85). Our data provides further evidence of a tight association between large transcriptional changes and successful stress-coping mechanisms.

### SCN transcriptomic profile

We observed significant upregulation of *Adora2a* in the SCN of resilient mice at ZT16. Adenosine is known to regulate *Per1* and *Per2* expression via A1/A2A receptor activation which is hypothesised to integrate light and sleep signalling to track homeostatic sleep pressure and regulate sleep architecture - both of which were previously found to be differentially impacted in resilient and susceptible mice (51,56,86). Upregulation of *Adora2a* in the SCN of resilient mice at ZT16 may be a subcomponent of adaptive homeostatic response that maintains consolidated sleep (51,56). Interestingly, the upregulation of A2A receptors in the forebrain is associated with depressive-like behaviour and decreased synaptic plasticity following unpredictable chronic mild stress (CMS) (87). Resilient mice displayed strong upregulation of *Ntrk1* and *Ngfr* in the SCN at ZT16. NGF binds to Ntrk1 and Ngfr (88,89). NGF levels are significantly lower in depressed patients (90) while increasing its levels has antidepressant properties (91,92). Furthermore, BDNF, which has functional roles in synaptic plasticity, stress response and antidepressant action in several regions, also binds to Ngfr (93,94). Ngfr is specifically synthesised in cholinergic neurons which strongly modulate SCN neuronal excitability and circadian rhythmicity at night (95–97).

We also observed high upregulation of cholinergic muscarinic receptors *Chrm 1-5.* These receptors have a functional role in fear and emotional responsiveness (98,99). Cholinergic signalling in the SCN can induce phase shifts in locomotor rhythms and activity (100). However, at present it is unclear how increased muscarinic receptor expression at ZT16 in the SCN drives resilience. A recent study demonstrated that the stress-associated neuropeptide CRH increases cholinergic interneuron firing that causes potentiation of dopamine transmission (101,102). Besides cholinergic receptors*, Drd2* and *Crh* were upregulated in the SCN of resilient mice at ZT16. Dopaminergic input from VTA to SCN modulates circadian entrainment through activation of excitatory Drd1 receptors (103). Upregulation of inhibitory *Drd2* in resilient mice might have some unknown homeostatic role in circadian regulation. Our observation that *Crh* is upregulated in resilient mice adds to the recent findings that CRH-expressing neurons are also present in the SCN (104). CRH increases gating of BDNF signalling (105). Thus, increased CRH expression at ZT16 may be responsible for putative BDNF-induced antidepressant-like effects in stress-resilient mice. The resilient phenotype has also been associated with increased cAMP and GPCR-mediated signalling (80,106). We found that these enriched pathways are predicted to be activated in the SCN of resilient mice. These are a subset of enriched plasticity-related pathways that are predicted to be activated in the SCN. However, the functional consequences of these changes are unclear at present since plasticity-related transcription factor *Creb3l1* was downregulated, while many others, such as *Gbx1*, were strongly upregulated in the resilient mice. Previously, *Creb* downregulation has been associated with stress resilience (107,108). Overall, few studies have examined stress effects on SCN physiology and those predominantly focused on the expression of clock genes (14,109–111). Our observations highlight potentially novel molecular mechanisms in the SCN that promote resilience to stress that warrant further investigation.

### Hb transcriptomic profile

*Adcy1* and *Adcy9* were upregulated in the Hb of resilient mice relative to controls at ZT16. Adenylyl cyclases play a pivotal role in cAMP-mediated and the ERK½-CREB signalling cascade and are known to modulate neuronal physiology (112–114). Overexpression of *Adcy1* in the forebrain prevented BDNF downregulation in stressed mice, thereby promoting stress resiliency (115). *Adcy1* forebrain overexpression induces greater long-term potentiation (LTP) (114). Moreover, we observed ZT16-specific upregulation of other plasticity-modulating genes, such as *Grin2a, Grin2b*, *Cacna1h*, *Camk2b*, and *Ppp3ca* which correlate with large predicted activation of synaptogenesis and LTP signalling pathways. Resilient mice exhibit greater hippocampal LTP relative to susceptible mice because of higher expression of *Grin2b* and *Camk2* (116). Upregulation of plasticity-related genes may be a part of a larger homeostatic response associated with stress resiliency (117,118). We also observed upregulation of *Gad1, Gabra5, Kcnq3* and *Npy1r* genes in stress-resilient mice. In contrast CMS impairs GABA release and uptake in the medial prefrontal cortex which is associated with depressive-like behaviour (119). Sub-chronic mild stress decreases long-term depression (LTD) of GABAergic neurons in the LHb (120). Furthermore, male mice with *Kcnq3* mutation that inhibits GABA binding exhibit reduced self-grooming, behaviour indicative of apathy (47,121). Neuropeptide Y action through its Y1 receptor has been associated with stress resilience and anxiolysis in several brain regions, including the Hb (122,123). In LHb, *Npy1r*-expressing neurons play a critical role in chronic stress adaptation by driving palatable food intake and reducing anxiety-like behaviour (124). Thus, it is possible that elevated *Npy1r* expression at ZT16 in the Hb drives increased food intake in resilient mice to compensate for lower levels of adipose tissue and serum leptin (125).

### Clock genes

Since our analysis revealed a strong effect of stress on the SCN and Hb at a specific timepoint, we examined the rhythmic expression of clock genes in both of these nuclei. Chronic stress induces arrhythmicity of *Arntl* in the SCN of both resilient and susceptible mice. Despite previous observation that knockdown of *Arntl* leads to behavioural despair, anxiety and helplessness in the CMS, our observation suggests that *Arntl* arrhythmicity in the SCN does not directly impact mood regulation (14). Another study found that CMS in rats induces perturbation of *Per2* in the LHb, without affecting the SCN rhythms (126). In our study, *Per1*, *Per3*, and *Cry1* were arrhythmic in the Hb of stress-naïve mice, but not resilient and susceptible mice. Although unexpected, these results can be attributed to the subdivisional differences at the level of MHb and LHb (127). Clock gene expression in the MHb has yet to be systematically investigated in mice, though it has been reported previously in hamsters and rats (128,129). Despite both the LHb and MHb showing diurnal variation in firing (35,43), the LHb displays more evident rhythms of higher amplitude than the MHb (13,42,127). In the present study we did not differentiate between the two subregions and reduced spatial resolution might have contributed to the observed arrhythmicity of clock genes in stress-naïve mice. Moreover, different stressors have been previously shown to differentially affect neural dynamics (84,130,131). Thus, the differential effect of CSDS and CMS on the transcriptomic landscape may be driven by type of stressor.

### Conclusion and future directions

The SCN and Hb are heterogeneous nuclei with many functionally and molecularly distinct subregions (69,132). Previous transcriptomic studies have examined only specific subpopulations of the Hb for purposes of circuit manipulations (26,133,134). Likewise, studies linking SCN and stress were limited to investigating a limited predetermined list of genes (14). To the best of our knowledge, no study has systematically examined the SCN or Hb bulk transcriptome in the context of murine chronic stress models across multiple timepoints throughout the circadian cycle. We are the first to show that the largest transcriptomic changes in the SCN and Hb of resilient mice occur at night. This highlights the importance of sampling over a 24-hour cycle to accurately determine dynamic changes in gene expression following stress exposure. Most studies fail to consider diurnal changes in expression profiles. Our time-dependent transcriptomic analysis suggests novel potential molecular mechanisms for driving resilience to stress that would not be evident within a constrained daytime sampling period. Such findings support time-dependent treatment administration that might enhance its efficacy. Crucially, our results highlight the link between temporal expression patterns and homeostatic stress responses.

## Acknowledgements

The authors have received funding from the following sources: NYUAD Start-Up Fund (DC), NYUAD Annual Research Fund (DC), NYUAD Research Enhancement Fund (DC), University Research Challenge Fund (DC), Al Jalila Research Foundation (AJF201638; DC), NYU Abu Dhabi Research Institute Award to the NYUAD Center for Genomics and Systems Biology (PN). The funders had no role in study design, data collection and analysis, decision to publish, or preparation of the manuscript.

We thank Justin Blau and Dan Ohtan Wang for critical reading of this manuscript.

After acceptance the RNA sequencing dataset will be available online at the Gene Expression Omnibus database.

## CRediT Statement

**Conceptualization:** Priyam Narain, Aleksa Petković, Marko Šušić, Dipesh Chaudhury

**Data curation:** Aleksa Petković, Marko Šušić. Priyam Narain

**Formal analysis:** Marko Šušić, Aleksa Petković, Priyam Narain, Nizar Drou

**Visualisation:** Marko Šušić, Aleksa Petković, Priyam Narain

**Validation:** Aleksa Petković, Priyam Narain, Marko Šušić

**Funding acquisition:** Dipesh Chaudhury

**Investigation:** Aleksa Petković, Marko Šušić, Priyam Narain, Salma Haniffa, Marc Arnoux, Mariam Anwar

**Methodology:** Priyam Narain, Aleksa Petković, Marko Šušić, Dipesh Chaudhury

**Project administration:** Priyam Narain, Dipesh Chaudhury

**Resources:** Dipesh Chaudhury

**Software:** Marko Šušić, Nizar Drou, Priyam Narain, Aleksa Petković

**Writing-original draft:** Aleksa Petković, Priyam Narain, Dipesh Chaudhury

**Writing-review-editing:** Aleksa Petković, Marko Šušić, Priyam Narain, Dipesh Chaudhury

**Supervision:** Priyam Narain, Dipesh Chaudhury

## Supplemental Figures

**Supplemental figure 1.**
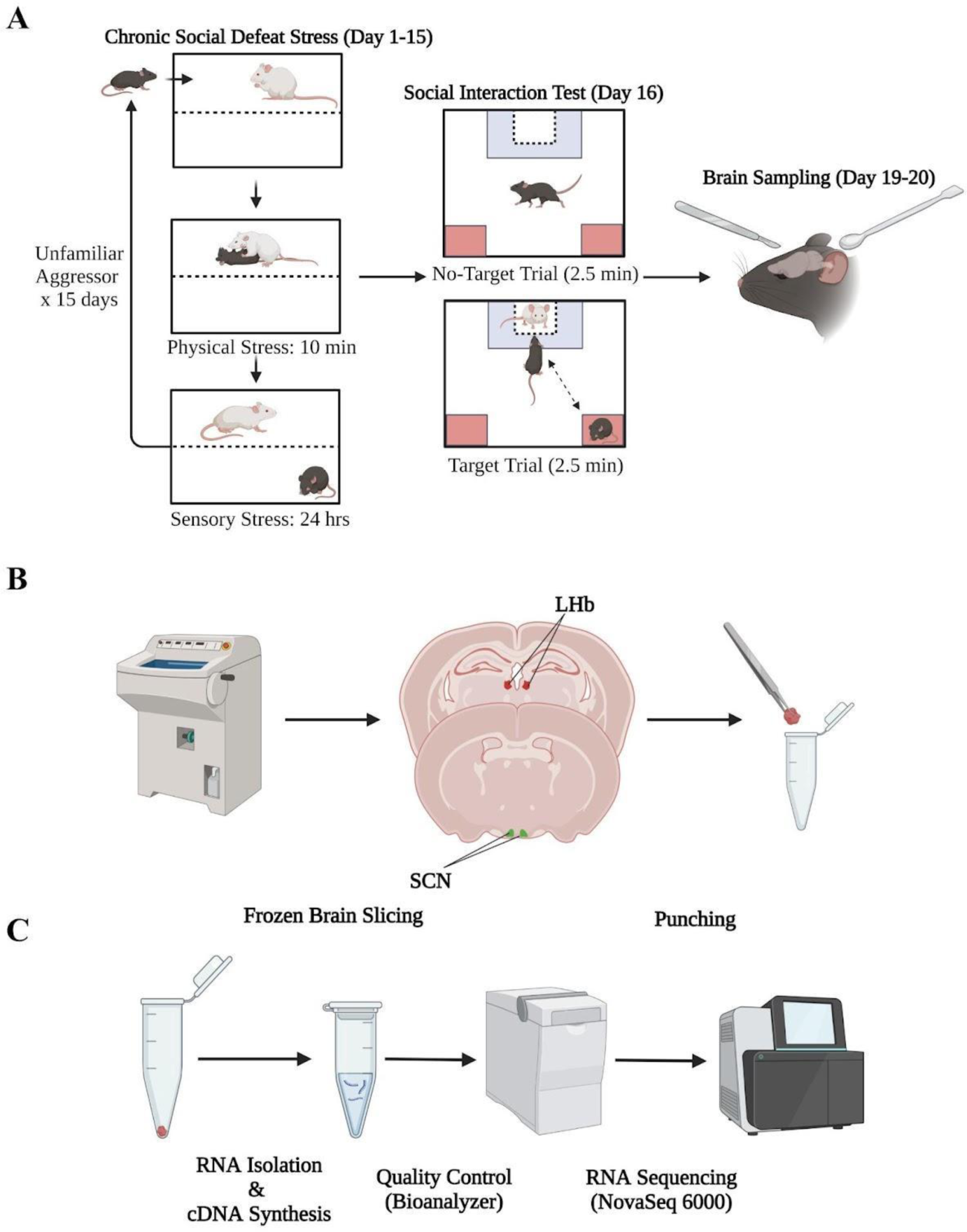
Graphical summary of the methodology. **(A)** Behavioural experimental methods and timeline. **(B)** Experimental methods for slicing and tissue punching. **(C)** Experimental methods for RNA isolation, quality control and sequencing.

**Supplemental Figure 2.**
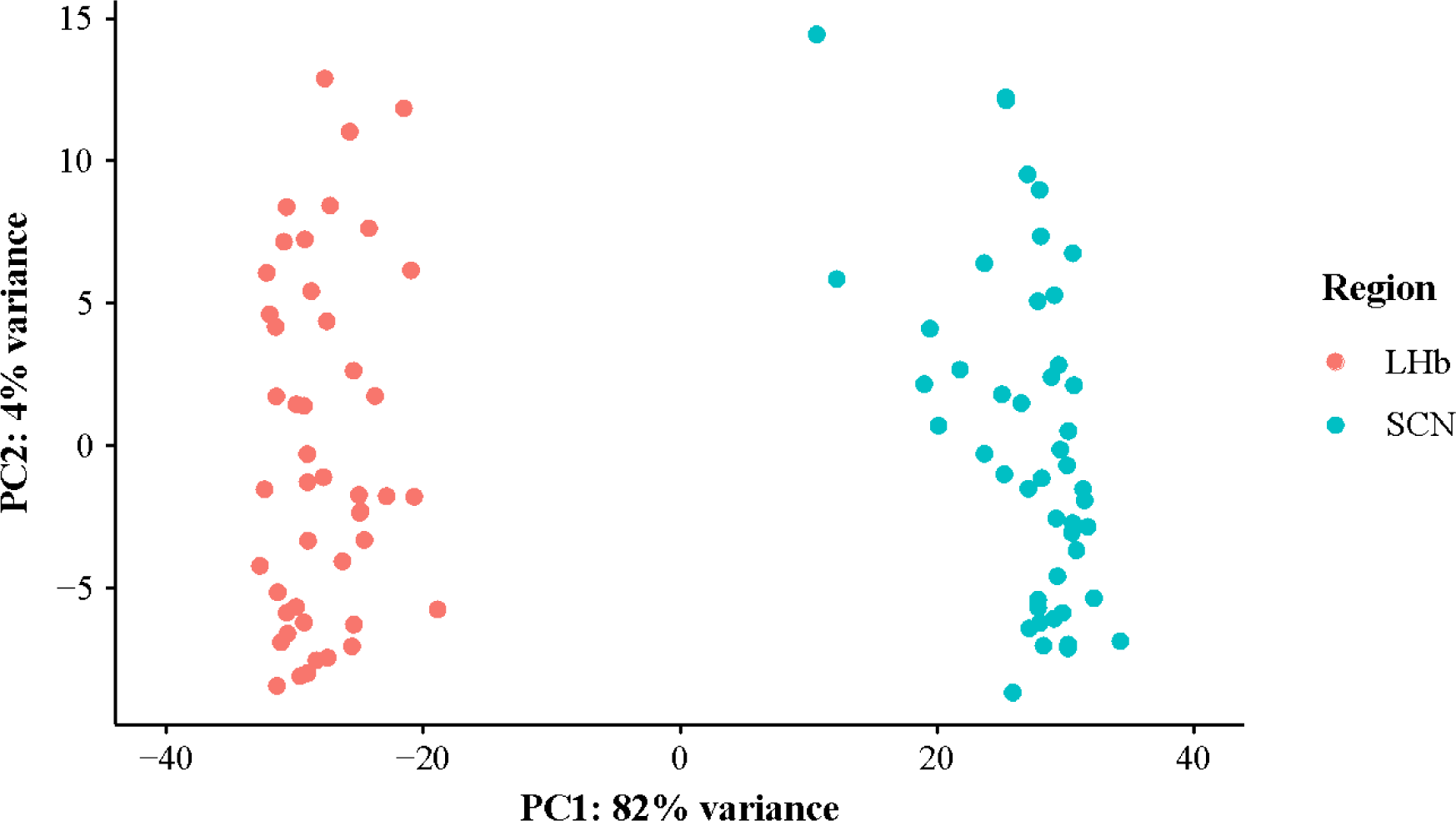
PCA plots showing clear separation between SCN and Hb samples. First principal component represents the regional identity of the samples and explains 82% of data variability.

**Supplemental Figure 3.**
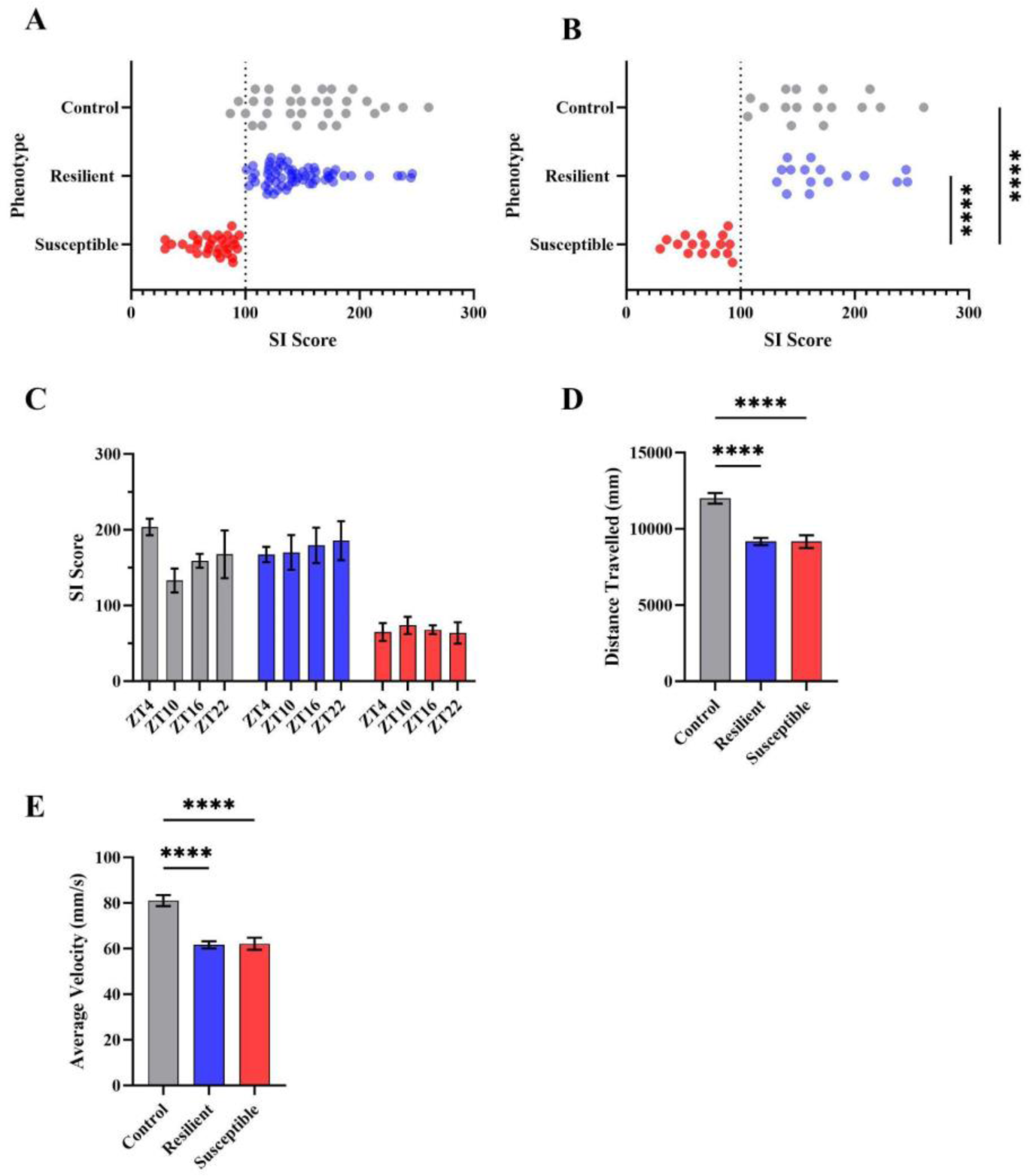
Behavioural measures. **(A)** One-way ANOVA revealed significant variation in SI scores (F(2,120) = 63.3, p < .001). Tukey’s post hoc comparison revealed significant differences between controls and susceptible mice (p < .001, 95% C.I. = [65.4, 106.8]), and resilient and susceptible mice (p < .001, 95% C.I. = [58.9, 94.9]). Stress-naïve and resilient mice did not differ in SI scores (p = .44, 95% C.I. = [-8.6, 26.9]). **(B)** One-way ANOVA revealed significant variation in SI scores of sequenced mice (F(2,45) = 45.3, p < .001). Tukey’s post hoc comparison revealed significant differences between sequenced control and susceptible mice (p < .001, 95% C.I. = [67.8, 128.6]), and resilient and susceptible mice (p < .001, 95% C.I. = [77.9, 138.4]). Stress-naïve and resilient mice did not differ in SI scores (p = .72, 95% C.I. = [-40.2, 20.6]). **(C)** Within phenotype, sample groups did not differ between timepoints in the SI scores. **(D)** One-way ANOVA for total distance travelled was significant (*F*(2,120) = 23.3, *p* < .001). The observed differences were significant between control and susceptible mice (*p* < .001, 95% C.I. = [1624, 4059]), and control and resilient mice (*p* < .001, 95% C.I. = [1796, 3889]) (Fig.1C). **(E)** One-way ANOVA for average velocity travelled was significant (*F*(2,120) = 24.2, *p* < .001). There were significant differences in average velocity between control and susceptible mice (*p* < .001, 95% C.I. = [10.8, 27]), and control and resilient mice (*p* < .001, 95% C.I. = [12.5, 26.4]). All values and bar plots are reported as mean±SEM. Significance levels: ** *p* < .01, *** *p* < .001.

